# Regulatory T cells restrain skin inflammation by modulating peripheral neuron activation

**DOI:** 10.1101/2024.05.14.594055

**Authors:** Alejandra Mendoza, Regina Bou-Puerto, Paolo Giovanelli, Stanislav Dikiy, Emma S. Andretta, Alexander Y. Rudensky

## Abstract

The skin integrates diverse signals discerned by sensory neurons and immune cells to elicit adaptive responses to a range of stresses. Considering interactions between nervous and immune systems, we questioned whether regulatory T cells (T_reg_ cells), a T cell subset that suppresses systemic and local inflammation, can modulate activation of peripheral neurons. Short-term ablation of T_reg_ cells increased neuronal activation to noxious stimuli independently from immunosuppressive function. We find that a population of skin T_reg_ cells is highly enriched for *Penk* expression, a precursor for endogenous opioid enkephalins. Acute depletion of Penk-expressing T_reg_ cells, or cell-specific ablation of *Penk* in T_reg_ cells increases neuronal activation in response to noxious stimuli and associated inflammation. Our study indicates that a population of T_reg_ cells exhibits neuromodulatory activity to restrain inflammation.

## Introduction

The gatekeeping function and integrity of barrier tissues are essential for organismal health and entail mitigation of harm elicited by noxious biotic and abiotic agents through proportionate, finely balanced responses. In the skin, diverse cell types communicate to enable integration of, and adequate reaction to a wide range of stimuli such as temperature, mechanical and chemical stresses, tissue damage, and infection. Cells of the nervous and immune systems sense potentially harmful deviations from normalcy and ensure the avoidance, clearance, or sequestration of the causative agents. The skin is endowed with a rich network of heterogeneous sensory nerve endings that initiate rapid protective behavioral and local tissue responses against diverse external stimuli by evoking sensations such as pain and itch(*1*). A growing body of work suggests that these neuronal cells can also expeditiously mobilize diverse innate and adaptive immune cell types and promote their effector functions through the release of neuromediators and other signaling molecules(*2-6*).

The regulatory T (T_reg_) cell lineage represents a principal specialized cellular component of the inhibitory arm of vertebrate immunity that promotes tolerance to ‘self’, commensal bacterial, dietary and environmental antigens, and limits responses to acute and chronic infections(*7*). At their non-lymphoid sites of residence, foremost barrier organs such as skin, T_reg_ cells undergo specialized adaptations to support tissue functions by modulating activity of both parenchymal and accessory cell types(*8-12*). Considering emerging realization of connectivity between nervous and immune systems, we questioned whether in barrier tissues T_reg_ cells can modulate proinflammatory neuronal activity by acting directly on peripheral neurons.

## Results

### Treg cells modulate peripheral neuron activation

To begin addressing this possibility we first examined potential physical interactions of T_reg_ cells with axons innervating the skin by proximity mapping **(Fig. 1a-b)**. We found that in the skin, T_reg_ cells closely juxtaposed axons, with 21.7% in direct contact (0 μm), 26.8% within 10 μm and 21.7% within 20 μm of the nearest axon **(Fig. 1c)**. This proximity raised the possibility of T_reg_ cell-neuron communications through direct cell-cell contacts or through short-range acting secreted factors. To assess potential effects of T_reg_ cells on sensory perception, we carried out short-term inactivation of T_reg_ cells and assessed its effect on neuronal activation by noxious stimuli(*5*). The primary somatosensory neurons that innervate the skin are pseudounipolar with their cell bodies housed in the dorsal root ganglia (DRG) and axons consisting of two branches: one that synapses at the dorsal horn of the spinal cord and another innervating the peripheral tissue(*13*). Monitoring expression of transcription factor cFos has been instrumental in identifying neural circuits, as its expression is induced rapidly after activation, and both cFos mRNA and protein have short half-lives in neurons(*14-16*). Using capsaicin, an activator of TRPV1 receptor expressed by nociceptive neurons, we assessed ability of T_reg_ cells to modulate the induction of cFos expression in the nuclei of neurons in the cervical DRG innervating the ear skin (*4, 17*)**(SFig.1a,b)**. Treatment of *Foxp3*^*DTR*^ mice with diphtheria toxin (DT) blocks protein synthesis in DTR expressing T_reg_ cells causing first their inactivation followed by death(*18-20*). At 18 hours post DT administration, we observed loss of function of skin T_reg_ cells as indicated by their impaired IL-10 production, while their numbers were largely unaffected **(SFig. 1c-f)**. This short-term DT treatment regimen allowed us to assess tempering of sensory neuron activity by skin T_reg_ cells without potential confounding effects of their death. Upon T_reg_ cell inactivation caused by DT, we observed heightened cFos induction in response to topical capsaicin application manifested in the increased percent of cFos^+^ nuclei (Tubβ3^+^) in cervical DRG **(Fig. 1d-e)**. The specificity of cFos staining in DRG was confirmed by using *Fos-gfp* reporter mice, where 92% of nuclei labeled by cFos antibody staining were also GFP positive **(SFig. 1g,h)** (*18*). Furthermore, short term DT treatment evoked a decrease in the behavioral response time to heat, indicating an increase in acute noxious thermal sensation in male mice **(Fig. 1f)**. Notably, this was not the case in female mice consistent with baseline sex-dependent differences in behavioral responses to thermal stimuli **(SFig. 1i)**(*21, 22*). However, the increased neuronal activation to noxious stimuli and sex-dependent differences in heat response were unlikely due to inflammation caused by T_reg_ cell inactivation, as we found no measurable increases in cytokine production or overall immune cell makeup of the skin after 18-hour DT treatment **(SFig. 1j-l)**. Furthermore, increased activation in DRG neurons was unlikely due to a direct response to DT, as mice lacking DTR did not show an increase in cFos^+^ Tubβ3^+^ nuclei in the DRG upon DT treatment (**SFig. 1m)**. These data suggest that T_reg_ cells prevent exacerbated neuronal activation in response to nociceptive stimulation.

**Figure 1.**
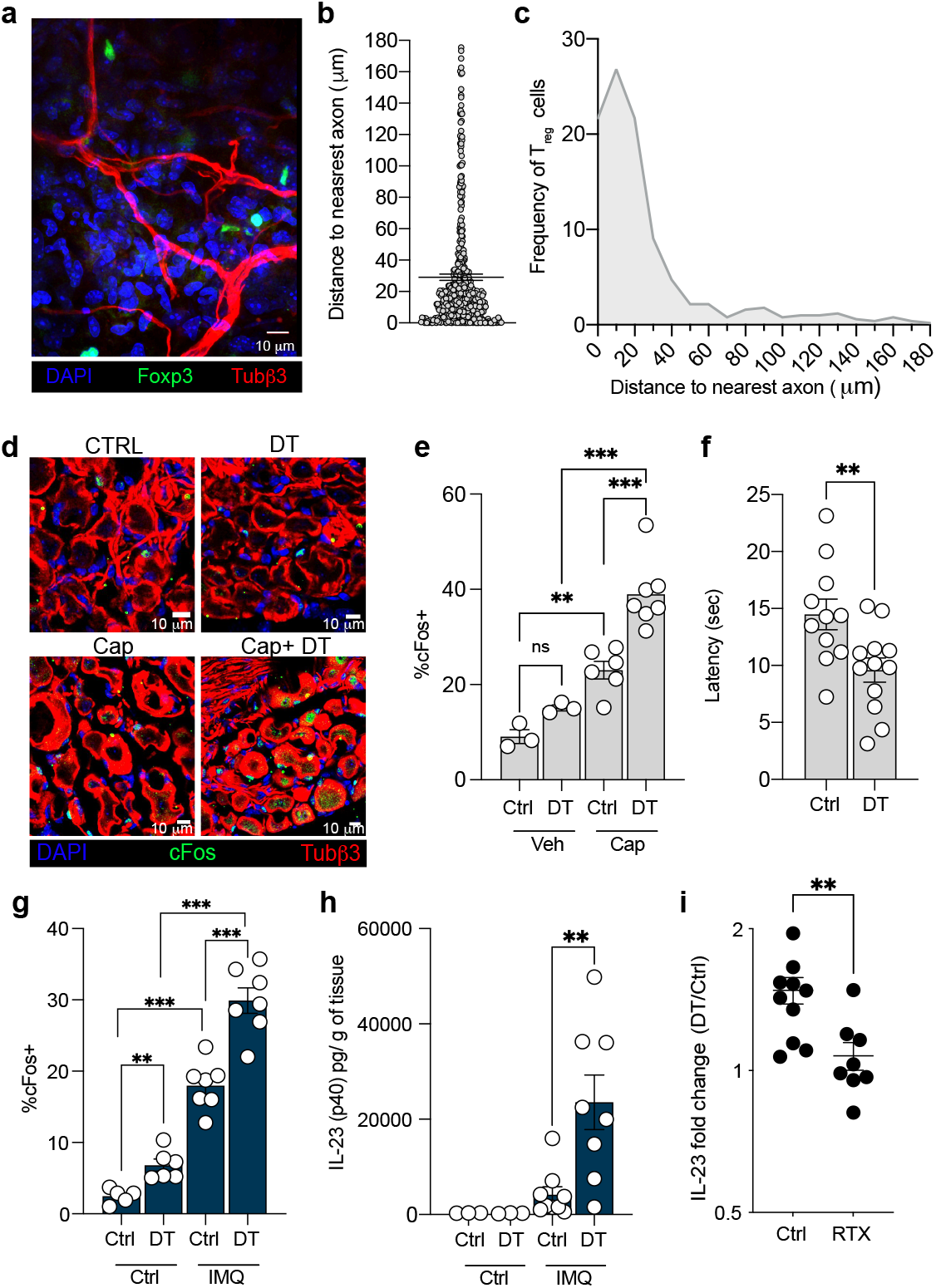
T_reg_ cell depletion increases nociception. **a-c**, Ear skin from *Foxp*^*gfp*^ mice was analyzed by confocal microscopy. **a**, Representative confocal microscopy image; sections were stained for Tubβ3 (red), DAPI (blue) and GFP (green). Scale bar, 10 μm. **b**, T_reg_ cell distance to nearest axon. Each dot represents a cell. Data compiled from ear skin of 3 mice. **c**, Histogram showing distribution of distances from T_reg_ cells to nearest axon. Data is shown as percentage of T_reg_ cells in 10 μm bins. Data compiled from ear skin of 3 mice. **d-f**, *Foxp3*^*DTR*^ mice were treated with DT or PBS control for 18 hours. **d-e**, Vehicle or capsaicin was applied to ear skin and DRG were isolated 30 min post-treatment. DRG were analyzed by confocal microscopy. **d**, Confocal microscopy images of DRG sections stained for Tubβ3 (red), DAPI (blue) and cFos (green). Scale bars, 10 μm. **e**, Percentage of cFos^+^ nuclei of neurons (Tubβ3^+^ cells) in DRG. Each dot represents average cFos^+^ nuclei from cervical DRG per mouse. **f**, Latency to respond to thermal stimulus was measured in male mice by hot plate assay. Each dot represents response time per mouse. **g-h**, *Foxp3*^*DTR*^ mice were treated daily with topical IMQ or petroleum jelly (control) on both ears (start: day 0). DT or PBS control was administered on day 1 and 2. Mice were analyzed on day 3. **g**, Percentage of cFos^+^ nuclei of Tubβ3^+^ cells in DRG. Each dot represents average cFos^+^ nuclei from cervical DRG per mouse. **h**, Ears were harvested, and total protein was prepared to quantify IL-23 (p40) per g of tissue by ELISA. **i**, *Foxp3*^*DTR*^ mice were treated with RTX or vehicle (DMSO). One week post RTX treatment mice were treated daily with IMQ (start: day 0). DT or PBS control was administered on day 1 and 2. Ears were collected on day 3 and IL-23 (p40) was measured by ELISA. IL-23 (p40) fold change between DT-treated and PBS-treated was calculated (DT/Ctrl) and compared between vehicle and RTX treated mice. Bars show means, error bars show SEM. **, p<0.001; ***, p<0.0001. P-values were calculated using an unpaired t-test.

### T_reg_ cells restrain exacerbated activation of peripheral neurons during early during inflammatory challenge

Cutaneous sensory neuron activation has been shown to be sufficient to trigger an inflammatory reflex arc that mobilizes immune effector cell function(*6*). To test if T_reg_ cell-neuron interactions in the skin have also an effect on inflammation-promoting neuronal activity, we used an established mouse model of psoriatic-like skin inflammatory response induced by the Toll like receptor 7 (TLR7) agonist Imiquimod (IMQ), where nociceptive sensory neurons have been shown to boost IL-23 production by dendritic cells in the initial phase of disease(*23*). To test if T_reg_ cells are required to restrict nociceptive neuron activation during the induction of psoriatic-like disease, we ablated T_reg_ cells in *Foxp3*^*DTR*^ mice 24 hours after IMQ challenge and analyzed mice on day 3 to allow for reliable detection of IL-23 levels. While DT induced T_reg_ cell depletion for the last 42 hours of IMQ challenge did not lead to increased T cell activation in the draining lymph nodes or CD4 T cell cytokine production **(SFig 1n-q)**, it did lead to increased neuronal activation in the DRG, reflected by higher frequency of nuclear cFos^+^ neurons in response to IMQ in comparison to control mice treated with PBS **(Fig. 1g)**. Consistent with increased neuronal activation, we found that ablation of T_reg_ cells resulted in heightened production of IL-23 **(Fig. 1h)**. This increase was blunted in mice pre-treated with resiniferatoxin (RTX), a toxin commonly used for selective TRPV1^+^ neuron targeting, which depletes TRPV1^+^ neurons alongside cold-sensing TRPM8 and TRPA1 neurons **(Fig. 1i**; **SFig. 1r-t)**. Thus, these results suggest that the increase in IL-23 production observed upon T_reg_ cell depletion in IMQ challenged *Foxp3*^*DTR*^ mice is dependent on sensory neurons. Overall, these data support the notion that T_reg_ cells restrain exacerbated activation of skin-innervating neurons during an inflammatory challenge thereby limiting skin inflammation during the early phase of psoriatic-like disease.

### A population of T_reg_ cells expresses *Penk*

To gain insights into potential mechanisms of the observed attenuation of activity of nociceptor sensory neurons we examined previously generated T_reg_ cell gene expression datasets for candidate genes, whose products have been implicated in signaling to peripheral nerves and modulating their function. This analysis identified *Penk*, a gene encoding the precursor neuropeptide proenkephalin, as a top candidate whose expression was highly upregulated in activated T_reg_ cells (*24-26*). Proenkephalin proteolytic cleavage results in the production of enkephalins, endogenous opioids known to inhibit pain perception by inducing analgesia(*27-29*). We hypothesized that a subset of T_reg_ cells expressing Penk negatively regulate sensory neuron activation to temper tissue inflammation in response to noxious stimuli. To examine the distribution, frequency, and localization of this subset of T_reg_ cells, we employed a historic reporter of Penk expression using mice expressing Cre recombinase driven by an IRES under the control of the endogenous *Penk* locus (*Penk*^*Cre*^) and a *R26-CAG-LSL-tdTomato* recombination reporter allele (*Rosa*^*LSL-tdTomato*^). Analysis of *Penk*^*Cre*^ *Rosa*^*LSL-TdTomato*^ mice showed tdTomato expression in a population of T_reg_ cells, characterized by its differential distribution among non-lymphoid organs with the highest prevalence in the skin but scarce in lymphoid organs and circulation **(Fig. 2a)**. Consistent with the fate-mapping results, we found a similar pattern of endogenous Penk transcript expression in spleen, lymph node and skin T_reg_ cells **(Fig. 2b)**. In support of potential crosstalk between *Penk* expressing (tdTomato^+^) T_reg_ cells and neurons we observed a relatively small, yet statistically significant enrichment of the former in the proximity of axons in the skin **(Fig.2c,d; SFig. 2a)**. Consistent with their specialized role, we found that Penk expressing T_reg_ cells remained highly prominent in the skin under inflammatory conditions triggered by IMQ, while their frequency stayed relatively low in the secondary lymphoid organs **(Fig. 2e)**. Notably, we observed Penk (tdTomato) expression by only minor proportion of conventional activated αβ T cells implying a specific role of Penk expressing T_reg_ cell subset in dampening nociceptive signaling **(SFig. 2b)**. To further examine features of skin Penk^+^ vs Penk^-^ T_reg_ cell subsets, we characterized their transcriptomes using plate-based SMART-seq2 (ss2) scRNA-seq analysis of skin T_reg_ cells isolated from IMQ challenged and control *Penk*^*Cre*^*R26-CAG-LSL-YFP* mice (*Rosa*^*LSL-YFP*^) **(SFig. 2c,d)**. Unsupervised clustering revealed three clusters of skin T_reg_ cells **(Fig. 2f)**. In both challenged and control mice, YFP expression was highly enriched in cluster 2, the cluster with the highest percent of cells with detectable Penk transcripts (**Fig. 2g; SFig. 2e**). Based on pseudotime analysis, cluster 2 represented the most differentiated cells amongst skin T_reg_ cells **(Fig. 2h)**. Furthermore, cells in cluster 2 were enriched for a T_reg_ cell specific TCR signaling and activation gene signature(*30*) **(SFig. 2f)** characterized by high expression of genes associated with T_reg_ activation, including *Tigit* and *Ctla4*, the transcription factor encoding genes *Hif1a, Nra4a1* and *Irf7*, as well as genes associated with migration, such as *Ccr8* and *Ccr2* **(Fig. 2i)**.

**Figure 2.**
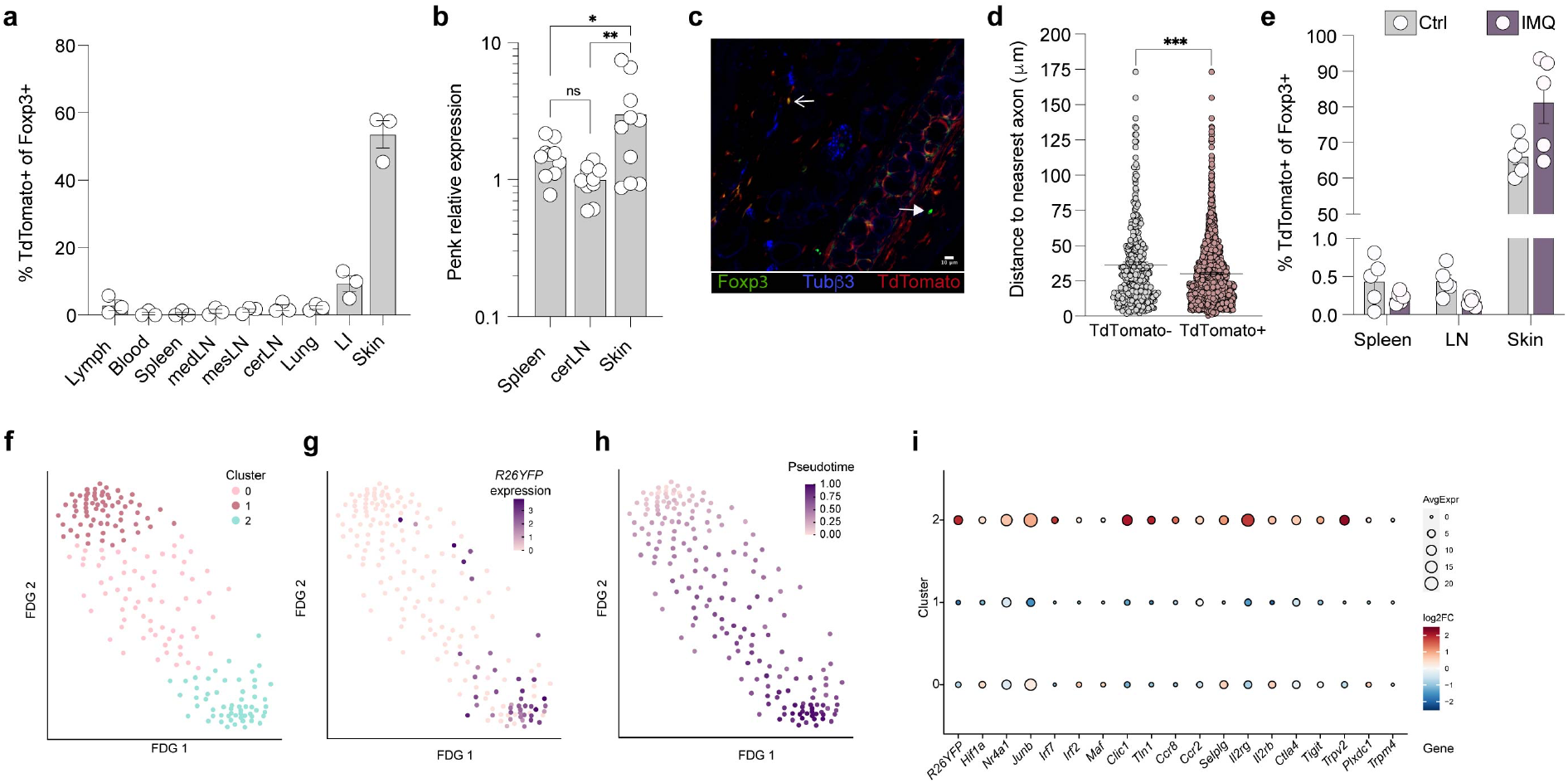
An effector population of T_reg_ cells express Penk. **a-e**, Distribution and localization of Penk-expressing T_reg_ cells in *Penk*^*Cre*^*R26*^*LSL-TdTomato*^ mice. **a**, Frequency of TdTomato^+^ T_reg_ cells in specified tissues from *Penk*^*Cre*^*R26*^*LSL-TdTomato*^*Foxp3*^*gfp*^ mice.**b**, RT-qPCR analysis of the expression of Penk in T_reg_ cells from spleen, skin-draining lymph node and skin. **c-d**, Ear skin from *Penk*^*Cre*^*R26*^*LSL-TdTomato*^*Foxp3*^*gfp*^ mice was analyzed by confocal microscopy. **c**, Representative confocal microscopy image of sections stained for Tubβ3 (blue), Foxp3-GFP (green) and tdTomato (red). Scale bar 10 μm. Open arrow shows tdTomato^+^ T_reg_ cell; closed arrow shows tdTomato^-^ T_reg_ cell. **d**, Distribution of distances from tdTomato^+^ and tdTomato^-^ T_reg_ cells to nearest axon in *Penk*^*Cre*^*R26*^*LSL-TdTomato*^*Foxp3*^*gfp*^ mice. **e**, *Penk*^*Cre*^*R26*^*LSL-TdTomato*^*Foxp3*^*gfp*^ mice were treated daily with topical IMQ or petroleum jelly (ctrl) on both ears for 3 days. Frequency of tdTomato^+^ T_reg_ cells in specified tissues. **f-i**, *Penk*^*Cre*^*R26*^*LSL-YFP*^*Foxp3*^*Thy1*.*1*^ mice were treated with petroleum jelly (control) or IMQ for 3 consecutive days. T_reg_ cells (Thy1.1^+^CD4^+^TCRβ^+^) were sorted from the ear skin of IMQ and control treated mice and processed for plate-based SMART-Seq2 scRNA-seq analysis. See Methods for details. **f**, Nearest-neighbor clustering was used to identify 3 distinct cell clusters. 2D force-directed graph layout of T_reg_ cells from IMQ or control treated ear skin, colored according to cluster. **g**, 2D force-directed graph layout of T_reg_ cell YFP expression. **h**, 2D force-directed graph layout of T_reg_ cells from IMQ or control treated ear skin. Shading (pink/low to purple/high) indicates cell pseudotime, as determined by the Palantir algorithm. See Methods for details. **i**, Plot depicting per cluster expression of select genes. Size of the points indicates the average expression per cell of each gene within each cluster, and color indicates log_2_ fold change of expression in that cluster versus all other cells. **a** and **d**, Dots represent data from individual mice, bars show means, error bars show SEM. **d**, Each dot represents distance from a single cell, lines show mean and error bars show SEM. Data were collected from 3 mice per group. *** p<0.001. P-values were calculated using an unpaired t-test. **f-i**, See methods for statistical analysis on SMART-Seq2 datasets.

### Penk expressing T_reg_ cells dampen peripheral neuron activation and inflammation

To directly test the possibility that Penk^+^ T_reg_ cells dampen nociceptive signaling, we sought to assess the effect of selective ablation of Penk^+^ T_reg_ cells on responses to noxious stimuli. First, we generated mice expressing *Penk*^*Cre*^ alongside *Gt(ROSA)26Sor* allele harboring the coding sequence for DTR preceded by a loxP site-flanked STOP cassette (*R26*^*iDTR*^)(*31*). In the resulting *Penk*^*Cre*^*R26*^*iDTR*^ mice all Penk expressing cells express DTR due to Cre-mediated excision of the STOP cassette rendering them susceptible to DT mediated ablation. Next, we generated bone marrow chimeras (Penk^Pos^DTR) by reconstituting irradiated αβT cell deficient *Tcrb*^*-/-*^ hosts with hematopoietic precursor cells from *Penk*^*Cre*^*R26*^*iDTR*^ and *Tcrb*^*-/-*^ mice mixed at a 1:4 ratio **(Fig. 3a**). This approach allowed ablation of Penk^+^ T_reg_ cells and a negligible population of effector αβT cells in Penk^Pos^DTR mice, while sparing other non αβ T cells that express Penk. DT-induced ablation of Penk^+^ T_reg_ cells in Penk^Pos^DTR mice had no effect on overall T_reg_ cell numbers in the lymph node. While ∼40% reduction T_reg_ cell numbers was observed in the skin, there were no measurable effects on effector T cells and their cytokine production or immune cell composition of the skin **(Fig. 3b-c; SFig. 3a-e)**. However, DT-induced ablation of Penk^+^ T_reg_ cells did increase sensitivity to noxious stimuli, manifested in shorter response times to heat at 18 hours after DT induced T_reg_ inactivation **(Fig. 3d)**. In addition, Penk^Pos^DTR mice showed increased induction of cFos in the DRG nuclei and expression of IL-23 in the skin in response to IMQ challenge when Penk^+^ cells when ablated with DT **(Fig. 3e-f)**. To complement these ‘loss-of-function’ experiments, i.e. Penk^+^ T_reg_ inactivation or ablation, we sought to perform a “retention-of-function” experiment by selectively retaining Penk^+^ T_reg_ cells while acutely depleting Penk^-^ T_reg_ cells, representing the vast majority of the overall Treg cell pool. To achieve that we took advantage of a *Foxp3*^*fl-DTR*^ allele, which contains a loxP site-flanked DTR coding sequence in the 3′UTR of the *Foxp3* gene(*32*). The latter was combined with *Penk*^*Cre*^ allele to generate Penk^Neg^DTR mice, in which Penk^+^ T_reg_ cells lose expression of DTR because its coding sequence is excised by Cre recombinase, whereas Penk^-^ T_reg_ cells retain DTR expression **(Fig. 3g)**. Despite a major systemic reduction in overall T_reg_ cell numbers, including the skin-draining lymph nodes (95% reduction), and a notable decrease in the skin (74% reduction), inactivation of Penk^-^ T_reg_ cells upon short term (18 hour) DT treatment of Penk^Neg^DTR mice did not affect heat sensitivity **(Fig. 3j**; **SFig. 3f-h)**. In contrast to depletion of Penk^+^ T_reg_ cells, cFos induction in DRG nuclei and skin IL-23 levels also did not change upon DT-induced Penk^-^ T_reg_ cell depletion under IMQ challenge **(Fig. 3k-l)**. Overall, mice with depletion of Penk^+^ T_reg_ cells had comparable fold increase in IL-23 to *Foxp3*^*DTR*^ mice, whereas the depletion of Penk^-^ T_reg_ cells showed no effect in comparison to controls **(Fig. 3m)**. Furthermore, the observation that the loss of Penk expressing T_reg_ cells can be numerically compensated for by non-expressing ones to a considerable extent, but not vice versa, supports the notion that Penk expressing T_reg_ cells represent a terminally differentiated effector population **(Fig. 2g; Fig 3c, i**). Together these data point to a specific role of Penk expressing T_reg_ cells in modulating nociceptive neuron activation and further suggest that the heightened basal response to noxious stimuli when depleting T_reg_ cells stems not from a systemic or local acute inflammatory response, but rather due to a paucity in a distinct function of a Penk-expressing T_reg_ cell subset separable from the immunosuppressive one. Consistent with this notion, we found that T_reg_ cell-specific ablation of IL-10, a potent immunosuppressive cytokine with a previously established role in T_reg_ cell mediated control of skin inflammation(*33*), had no effect on thermal sensitivity, or on the level of cFos induction in DRG neurons, or on IL-23 production in the early response to IMQ treatment **(SFig. 4a-e)**.

**Figure 3.**
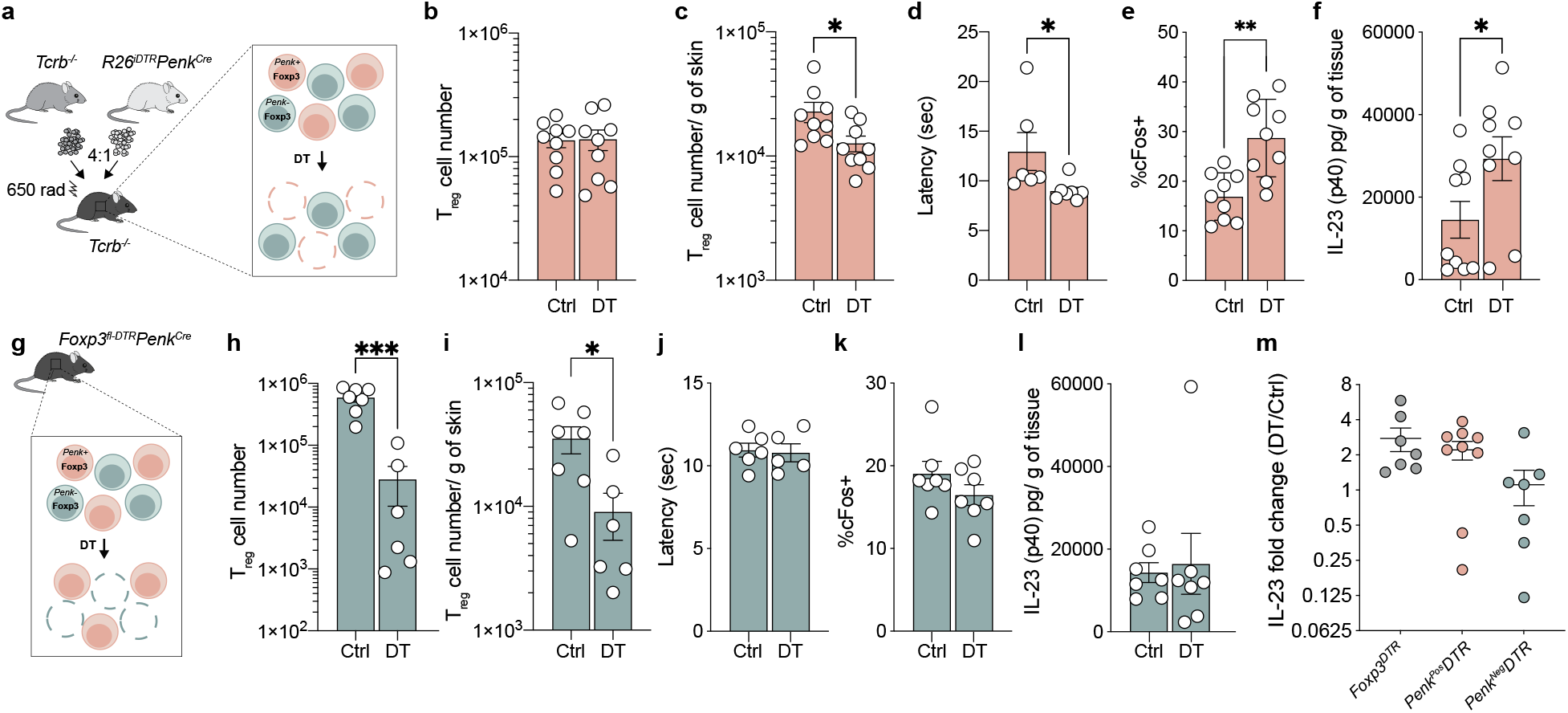
Acute ablation of Penk-expressing T_reg_ cells results in increased nociception and inflammation. **a-f**, *Tcrb*^*-/-*^ mice were irradiated and reconstituted with a mix of bone marrow from *Tcrb*^*-/-*^ and *R26*^iDTR^*Penk*^*Cre*^ at 4:1 ratio (referred to as *Penk*^*pos*^*DTR*). 6 weeks post reconstitution *Penk*^*pos*^*DTR* mice were treated with IMQ for 3 consecutive days (start: day 0). DT or PBS control was administered on day 1 and 2. Tissues were collected on day 3. **a**, Schematic for bone marrow chimeric mice and depletion of Penk-expressing T_reg_ cells. **b-c**, Number of T_reg_ cells found in the cervical lymph nodes (b) and skin (c) following DT treatment in *Penk*^*pos*^*DTR* mice. **d**, Latency to respond to thermal stimulus was measured in male *Penk*^*pos*^*DTR* mice by hot plate assay 18 hours post DT or PBS control treatment. **e**, Percentage of cFos+ nuclei of Tubβ3 + cells in DRG in IMQ treated *Penk*^*pos*^*DTR* mice. **f**, Ear skin from *Penk*^*pos*^*DTR* mice were harvested, and total protein was prepared to quantify IL-23 (p40)/ g of tissue by ELISA. **g-l**, *Foxp3*^fl-DTR^*Penk*^*Cre*^ mice (referred to as *Penk*^*neg*^*DTR*) were treated with IMQ for 3 consecutive days (start: day 0). DT or PBS control was administered on day 1 and 2. Tissues were collected on day 3. **g**, Schematic of depletion of non-expressing *Penk* T_reg_ cells in *Penk*^*neg*^*DTR* mice. **h-i**, Number of T_reg_ cells found in the cervical lymph nodes (h) and skin (i) following DT treatment in *Penk*^*neg*^*DTR* mice. **j**, Latency to respond to thermal stimulus was measured in male *Penk*^*neg*^*DTR* mice by hot plate assay 18 hours post DT or PBS control treatment. **k**, Percentage of cFos+ nuclei of Tubβ3 + cells in DRG in IMQ treated *Penk*^*neg*^*DTR* mice. **l**, Ears from *Penk*^*neg*^*DTR* mice were harvested, and total protein was prepared to quantify IL-23 (p40)/ g of tissue by ELISA. **m**, Comparison of IL-23 fold change DT treated and PBS treated (DT/Ctrl) *Foxp3*^*DTR*^, *Penk*^*pos*^*DTR* and *Penk*^*neg*^*DTR* mice following daily IMQ treatment (start: day 0) followed by DT administration on day 1 and 2. Each dot represents data from a mouse, bars show mean, error bars show SEM. *, p<0.005; **, p<0.001; ***, p<0.0001. P-values were calculated using an unpaired t-test.

### *Penk* expression in T_reg_ cells restrains peripheral neuron activation and cutaneous inflammation

The observed IL-10 independent regulation of nociception by a T_reg_ cell subset expressing Penk suggested that this particular function can be at least partially dependent on the production of enkephalins. Gene expression analysis showed T_reg_ cell expression of processing enzymes capable of proenenkephalin proteolytic cleavage (*Pcsk1, Ctsl* and *Furin*)(*34-37*). Signaling through opioid receptors by enkephalins modulates pre- and postsynaptic calcium channels, leading to attenuation of neuron excitability and reduction of neuropeptide release in peptidergic neurons(*38*). In addition, opioid receptor activation can prevent neuronal excitation and propagation of action potentials(*39-45*). To test the function of T_reg_ cell derived enkephalins, we generated *Foxp3*^*creERT2*^ mice expressing a conditional *Penk* allele (*Penk*^*fl*^) and ablated *Penk* gene expression in T_reg_ cells using tamoxifen **(SFig. 5a-c)** (*46*). Extended deletion of *Penk* (28 days) in T_reg_ cells produced a mild increase in cFos^+^ nuclei in the DRG, without any signs of increased immune activation or inflammation at major barrier sites, including the skin and intestine, or in secondary lymphoid organs compared to littermate *Penk*^*fl/wt*^*Foxp3*^*creERT2*^ controls **(SFig. 5d-q**). However, consistent with a minor but significant increase in basal neuronal activation in the DRG, *Penk*^*fl/fl*^*Foxp3*^*creERT2*^ mice exhibited increased sensitivity to heat as their response time to thermal stimulus decreased compared to littermate controls **(Fig. 4a)**. To test the role of T_reg_ cell derived enkephalins under inflammatory conditions *Penk*^*fl/fl*^*Foxp3*^*creERT2*^ and littermate control mice were treated with tamoxifen and challenged with IMQ one week later. *Penk* ablation in T_reg_ cells resulted in higher frequencies of cFos^+^ neurons in the DRG compared to littermate controls in response to IMQ **(Fig. 4b)**. The heightened neuronal activation was accompanied by increased IL-23 production in the early phase of disease **(Fig 4c)**. IL-23 has been shown to act on IL-23R^+^ dermal γδ T cells to induce IL-17 production, which facilitates the recruitment of circulating neutrophils in psoriasiform skin inflammation(*23*). Consistent with higher neuronal activation in the DRG and IL-23 levels, we found increased frequency of IL-17 producing dermal γδ T cells, and increased numbers of neutrophils in *Penk*^*fl/fl*^*Foxp3*^*creERT2*^ vs. littermate control mice after 7 days of IMQ challenge (**Fig. 4d,e**; **SFig. 6a-e)**. This was accompanied by increased ear swelling compared to controls and increased thickening of the epidermal layer, hallmarks of IMQ-induced inflammation **(Fig 4f,g)**. As enkephalins act upon members of the opioid receptor family, we examined published single cell RNA-seq datasets for skin cells and DRG and observed their expression exclusively in neuronal populations, suggesting that Penk expressing T_reg_ cells are acting directly on neurons(*47-50*). A corollary of this assumption is that the function of Penk^+^ T_reg_ cells in regulation of early phase skin inflammation were dispensable in the absence of nociceptive capacity. To experimentally test this possibility, we ablated nociceptors in *Penk*^*fl/fl*^*Foxp3*^*creERT2*^ mice and littermate controls by RTX administration **(SFig. 6f,g)**. RTX treated *Penk*^*fl/fl*^*Foxp3*^*creERT2*^ mice had no increase in IL-23 production compared to littermate controls in response to IMQ in contrast to mice that had not undergone sensory denervation (**Fig. 4h, SFig. 6h**). These results suggest that the effects of Penk deficiency in T_reg_ cells on inflammation were downstream of sensory neurons as ablation of TRPV1^+^ sensory nerves negated the effect of T_reg_ cell specific *Penk* deficiency.

**Figure 4.**
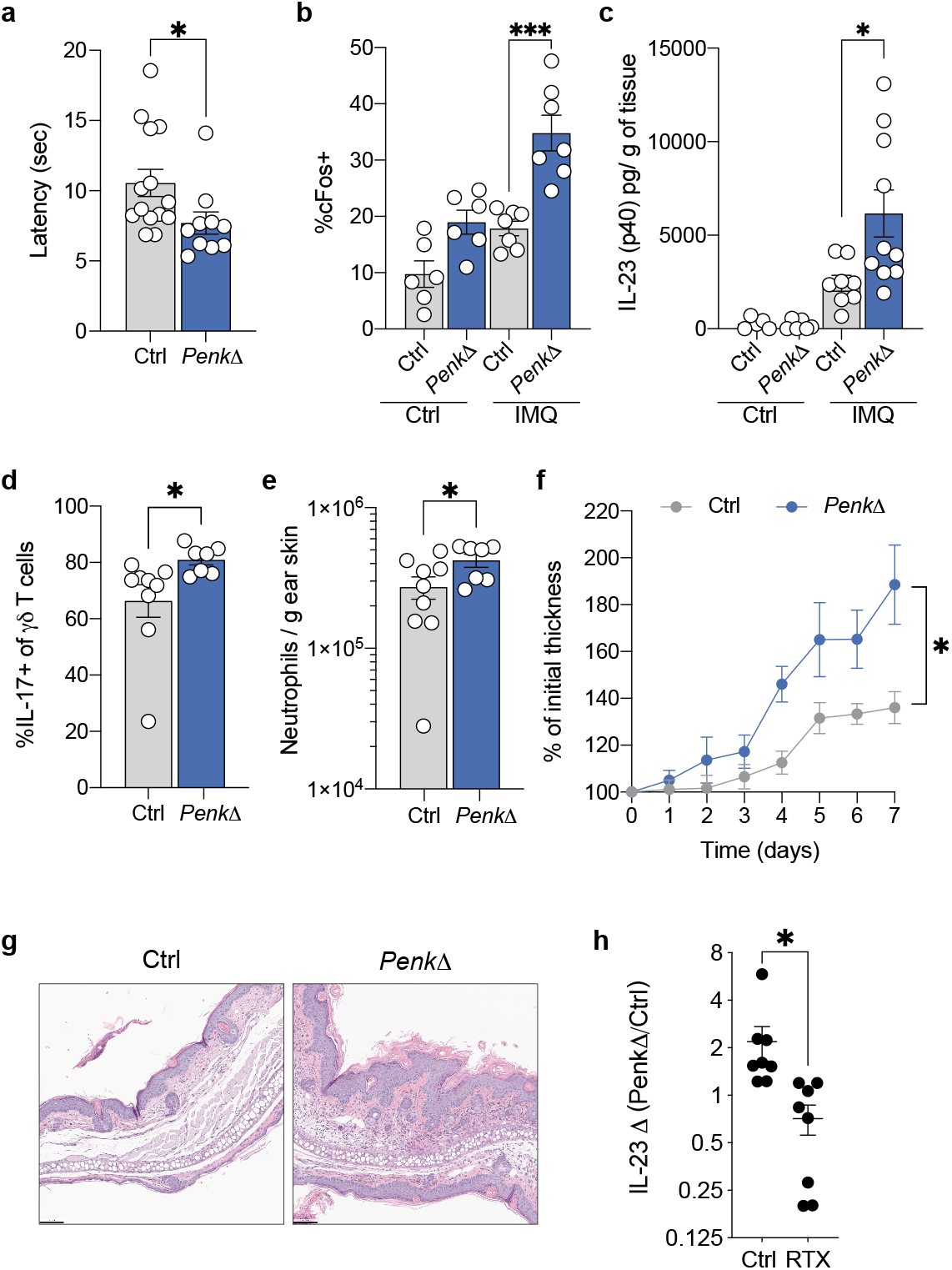
T_reg_ cell *Penk* expression is required to prevent exacerbated nociception and inflammation. **a-c**, *Penk*^*fl/wt*^*Foxp3*^*creERT2*^ (Ctrl) and *Penk*^*fl/fl*^*Foxp3*^*creERT2*^ (*PenkΔ*) were treated with 2 doses of tamoxifen, 3 days apart. **a**, Latency to respond to thermal stimulus was measured in male Ctrl *and PenkΔ* mice by hot plate assay 7 days post tamoxifen administration. **b-c**, 7 days post tamoxifen administration male Ctrl and *PenkΔ* mice were treated daily with IMQ or control (petroleum jelly) for 3 days. **b**, Percentage of cFos^+^ nuclei of DRG neurons (Tubβ3^+^ cells). **c**, Concentration of IL-23 (p40)/ g of ear tissue quantified by ELISA. **d-g**, *Penk*^*fl/wt*^*Foxp3*^*creERT2*^ (Ctrl) and *Penk*^*fl/fl*^*Foxp3*^*creERT2*^ (*PenkΔ*) were treated with 2 doses of tamoxifen, 3 days apart. 7 days post tamoxifen administration male Ctrl and *PenkΔ* mice were treated daily with IMQ for 7 days. **d**, Percentages of dermal γδ T cells producing IL-17A 3 hours post restimulation with PMA/ionomycin from Ctrl and *PenkΔ* mice. **e**, Total number of neutrophils in skin tissue from Ctrl and *PenkΔ* mice. **f**, Ear thickness was measured at indicated time points, data shows percent of thickness prior to IMQ treatment (n=7–8 mice per time point). **g**, Representative histological sections of IMQ treated ears at day 7 stained by H&E. **h**, *Penk*^*fl/wt*^*Foxp3*^*creERT2*^ (Ctrl) and *Penk*^*fl/fl*^*Foxp3*^*creERT2*^ (*PenkΔ*) mice were treated with RTX or vehicle (DMSO). One week post RTX treatment mice were treated with 2 doses of tamoxifen 2 days apart. 7 days following the last tamoxifen dose mice were treated daily with IMQ (start: day 0). Ears were collected on day 3 and IL-23 (p40) was measured by ELISA. IL-23 (p40) fold change between *PenkΔ* and Ctrl was calculated (*PenkΔ* /Ctrl) and compared between vehicle and RTX treated mice. Each dot represents data from a mouse, bars show mean, error bars show SEM. *, p<0.005; ***, p<0.0001. P-values were calculated using an unpaired t-test.

## Discussion

In the skin, a barrier tissue exposed to a wide range of external stimuli, extensive cell-cell communications relay signals from sensors to effectors invoking both positive and negative regulation of their activity. Through activation of these homeostatic cellular circuits, sensory neurons play a key role in the modulation of local immune cell activity and in preservation of barrier tissue integrity and function. We identified T_reg_ cells as a prominent modulatory component of this neuro-immune circuitry that prevents exacerbated activation of peripheral neurons and thereby tempers skin inflammation. Our results indicate that a population of T_reg_ cells acts to oppose exacerbated activation of sensory neurons through the production of an endogenous opioid that reduces nociceptive signaling and by limiting its tone, to restrain associated inflammation during inflammatory challenge. Previous studies of germline ablation of *Penk* gene as well as opioid receptors strongly suggest that tonic endogenous opioid signaling is required for preventing exacerbated nociception, likely at peripheral sites as well as within the brain and spinal cord(*27, 51*). These studies are in line with our observations that enkephalin production from skin T_reg_ cells tones down nociceptor neuron sensitivity to noxious stimuli thereby preventing exacerbated inflammation. Pain is a hallmark of injury, inflammation, infection, and neuropathy. As key components of the suppressive arm of immunity, T_reg_ cells have been anticipated to decrease sensory neuron activation solely by reducing inflammation. While T_reg_ cells may prevent the sensitization of nociceptors indirectly by reducing the local abundance of pro-inflammatory mediators, our studies suggest that they can also directly act on peripheral neurons to reduce nociception. Thus, the production of enkephalins represent a novel neuromodulating modality of T_reg_ cells required to prevent exacerbated sensory responses to noxious stimuli required to dampen inflammation. While blocking peripheral neuron responses to noxious stimuli would render the organism vulnerable to environmental insults, the converse - exaggerated responses - can result in the spurious development of maladaptive states. In this context, our studies suggest that T_reg_ cells play a non-redundant role in moderating the tone of responses to noxious insults by peripheral neurons.

## Acknowledgments

We thank A. Zimmer for generating and providing *Penk*^*Fl*^ mice, A. Bravo for help with animal husbandry, and all the members of the Rudensky laboratory for technical input and discussion. This work was supported by the National Institutes of Health/National Cancer Institute Cancer Center Support Grant P30 CA008748 (A.Y.R.), National Institutes of Health grant R01 AI034206 (A.Y.R.), Bristol Myers Squibb Fellowship from the Cancer Research Institute (A.M.), Basic and Translational Immunology Postdoctoral Award from Ludwig Cancer Research Center, Memorial Sloan Kettering Cancer Center (A.M.), Postdoctoral Fellowship from the Center for Experimental Immuno-oncology, Memorial Sloan Kettering Cancer Center (A.M.), National Institutes of Health grant T32 CA009149 (A.M.), Kravits WiSE Graduate Fellowship (R.B.P.), Dorris J. Hutchison Pre-Doctoral Fellowship (P.G.), and the Ludwig Center at the Memorial Sloan Kettering Cancer Center. A.Y.R. is an investigator with the Howard Hughes Medical Institute.

## Author contributions

A.M. and A.Y.R. designed the experiments. A.M., R.B.P., P.G., S.D., and E.A. performed experiments and analyzed data. A.M. and A.Y.R. interpreted the data and wrote the manuscript.

## Data availability

Raw and processed sequencing data (smart-seq2) will be made available through the GEO database (Accession number pending). The codes used for image analysis can provided upon request.

## Supplementary Materials

Materials and Methods

References (*52–74*)

Supplementary Figures S1 to S6

