## Supplementary materials for "Regulatory T cells restrain skin inflammation by modulating peripheral neuron activation"

**The PDF file includes:**

Materials and Methods
References (*52–74*)
Supplementary Figures S1 to S6

### Materials and Methods

#### Mice

*Foxp3*$^{gfp}$, *Foxp3*$^{DTR}$, *Foxp3*$^{Thy1.1}$, *Rosa26*$^{LSLTdtomato}$, *Rosa26*$^{LSLYFP}$, *R26*$^{iDTR}$, *Foxp3*$^{fl\text{-}DTR}$, *Il10*$^{fl}$, *Tcrb*$^{-/-}$, *Fos-gfp and Foxp3*$^{creERT2}$ mice were described previously(*52-61*). *Penk*$^{Cre}$ mice were purchased from Jackson Laboratories (stock ID: 025112). Frozen *Penk*$^{fl}$ sperm was generously provided by Andreas Zimmer. Treatments of mice were performed under protocol 08-10-023 approved by the Sloan Kettering Institute (SKI) Institutional Animal Care and Use Committee. All mice were analyzed at 8-12 weeks, except for bone marrow chimeras which were analyzed at 12-15 weeks of age. All experiments show a mix of females and males with sex-matched littermate controls except for behavioral test to heat. All mouse strains were maintained in the SKI animal facility in specific pathogen free (SPF) conditions in accordance with institutional guidelines and ethical regulations.

#### Diphtheria toxin treatment

Mice were injected i.p. with 1.25 μg diphtheria toxin (DT) (List Biological Laboratories) in PBS and analyzed 18 hours after a single dose or two daily consecutive doses.

#### Capsaicin treatment

*Foxp3*$^{DTR}$ mice were treated with DT. 18 hours following DT treatment mice were anesthetized with 2.5% isofluorane and 250 μg capsaicin was applied topically to dorsal and ventral sides of ear skin in acetone and olive oil solution (3:1). Mice were analyzed 30 minutes after capsaicin application.

#### Heat threshold testing

Noxious thermal sensation in mice of indicated genotypes or pharmacological treatments was measured by hot plate assay or tail flick tests. For hot plate test, mice were placed on a surface maintained at 55-56°C. The time for mice to hindpaw lick and/or shake, or jump, was recorded, and mice were immediately removed from the hot plate (animals were kept in hot plate for a maximum of 25 seconds). For tail flick test, mice were placed on acrylic mouse restrainer. A water bath maintained at 48 °C was used to immerse the distal 2/3 of each mouse's tail. The latency until withdrawal (rapid flick) was measured and tails were immediately removed from water (tails were kept in water for a maximum of 25 seconds).

#### Imiquimod treatment

Mice of indicated genotypes or pharmacological treatments were anesthetized with 2.5% isofluorane and treated with 5% Imiquimod (Perrigo) applied topically to dorsal and ventral sides of ear skin. Mice were treated starting on day 0 three times and euthanized for analyses on day 3 or were treated seven times and euthanized for analyses on day 7. Ear thickness was measured using a Kafer Thickness Gauge (J15, Long Island Indicator Service).

**Sensory denervation**

Resiniferatoxin (RTX), a capsaicin analogue, was injected subcutaneously into the flank of 6-week-old mice in three escalating doses (30 µg /kg, 70 µg /kg and 100 µg /kg of RTX purchased from Abcam, or 10 µg /kg, 25 µg /kg and 50 µg /kg of RTX purchased from AdipoGen Life Sciences) on consecutive days. Control mice were treated with vehicle (DMSO in PBS). Denervation was confirmed a week following treatment by tail flick or hot plate assay.

**Generation of bone marrow chimeric mice**

*Tcrb*$^{-/-}$ recipient mice were irradiated (650 Gy). The following day, bone marrow was isolated from femurs and tibias of donor mice. $2\times10^6$ total bone marrow cells were transferred into recipient mice via retro-orbital injection. Mice were analyzed 6 weeks after bone marrow reconstitution.

**Tamoxifen administration**

40 mg/ml tamoxifen was dissolved in corn oil (Sigma). Mice were orally gavaged with two doses of 200µl tamoxifen emulsion administered in 3 days. Mice were treated with Imiquimod 1 week following the last tamoxifen dose. For long term tamoxifen treatment, mice were orally gavaged with a dose of 200µl tamoxifen emulsion administered once per week.

**RT-qPCR**

Flat bottom 96-well plates were coated with anti-CD3/CD28 or anti-CD28 alone at 1 µg/mL in PBS, overnight at 4°C. $T_{reg}$ cells were sorted from *Foxp3* reporter mice from secondary lymphoid organs and plated in anti-CD3/CD28 coated plates at $10^5$ cells per well in 150µL complete RPMI with 1000U/ml of IL-2. For glucocorticoid stimulation $T_{reg}$ cells were treated with dexamethasone (Sigma-Aldrich) at the indicated concentrations for 24 hours at 37°C. Total RNA from $T_{reg}$ cells was isolated using RNeasy Plus Micro Kit (QIAGEN) and cDNA was synthesized using qScpript cDNA Synthesis Kit (QuantaBio) according to manufacturers' instructions. *Penk* transcript levels were quantified via RT-qPCR, using PowerUp SYBR Green Master Mix and the QuantStudio 6 Flex Real-Time PCR System (Applied Biosystems). Expression levels were determined with the

$2^{-\Delta\Delta CT}$ method, using Actb expression as an internal control. Penk primers: GAGAGCACCAACAATGACGAA (forward) and TCTTCTGGTAGTCCATCCACC (reverse); Actb primers: CTCAGGAGGAGCAATGATCTTGAT (forward) and ACCACCATGTACCCAGGCA (reverse).

**Retrograde tracer labelling**

Intradermal injection into the ear skin was performed using a 31G needle for retrograde tracing. A total of 10 µl of 0.1% CTB-488 (Invitrogen) in PBS was injected into each ear intradermally. DRG tissues were taken 5 days after injection to allow the dye to reach the DRG soma. DRG were analyzed by microscopy or digested for flow cytometry.

**Microscopy**

Confocal imaging was done using standard conditions. In brief, ear skin was excised, fixed for 2 hours at room temperature in 4% paraformaldehyde. For imaging DRG, mice were euthanized and cervical region of the spine was immediately fixed in 4% paraformaldehyde overnight at 4°C. Fixed DRG were isolated from spine. Following fixation, tissues were dehydrated at 4°C in 30% sucrose in PBS. Tissues were snap frozen in OCT compound (Sakura Tissue-Tek). 100 µm ear skin sections were cut and stained free-floating. DRG tissue was cut into 10 µm sections, slides were then fixed in acetone for 20 minutes at -20°C and dried overnight. Tissue sections were rehydrated in PBS for 10 minutes prior to staining. Staining was done in 10% donkey serum (Jackson ImmunoResearch) 0.3% Triton X-100 in 1x PBS overnight for skin and 24 hours for DRG. For quantification of cFOS in Tubβ3+ cells in DRG, images of 6-10 cervical DRG per mouse were taken, quantified and averaged. Tissues were imaged in Slow Fade mounting media (Life Technologies). All images were acquired using an SP8 (Leica) confocal microscope with a 40X oil immersion objective. Images were processed and analyzed using ImageJ software (version 2.0.0-rc-54/1.51h; National Institutes of Health). The codes used for image analysis can be provided upon request.

**Cell isolation and flow cytometry**

To label intravascular cells, mice were injected intravenously with 3 µg of anti-CD45 antibody. 3 minutes after injection mice were euthanized. Blood was collected from the vena cava and red blood cells were lysed in 1x ACK lysis buffer (155mM Ammonium Chloride, 10mM Potassium Bicarbonate, 100nM EDTA pH 7.2) for 3 minutes at room temperature prior to staining. Lymph was collected from the cisterna chyli into acid citrate-dextrose solution (Sigma). Lymphocytes were isolated from lymph nodes and spleen by mechanical disruption and filtration through a 100 μm cell strainer in Staining buffer (2%FBS, 1mM EDTA, 10 mM HEPES, 1x PBS). Ear skin tissue was weighed, separated into dorsal and ventral sides, and placed in 5mL snap-cap tubes (Eppendorf) in 3mL Wash buffer (1x RPMI 1640 w/ 2% FBS, 10mM HEPES buffer, 1% penicillin/streptomycin, 2mM L-glutamine) supplemented with 0.2U/mL collagenase A (Sigma) and 1U/mL DNase I (Sigma), along with three ¼ inch ceramic beads and shaken horizontally at 250RPM for 25 minutes at 37°C. Supernatant was removed and kept on ice while remaining undigested tissue was placed back on collagenase A and DNase I containing media and ceramic beads for an additional 20 minutes. Digested skin samples were combined and passed through a 100 μm strainer and centrifuged to remove collagenase solution. Large intestine was measured, defatted, opened longitudinally and luminal contents were removed by vigorous manual shaking of the tissue in PBS. DRG were digested in 1 ml of Wash buffer supplemented with 0.2U/mL collagenase A (Sigma) and 1U/mL DNase I (Sigma), in 2 ml tube along with one ¼ inch ceramic bead and shaken horizontally at 250RPM for 20 minutes at 37°C. Digested DRG suspensions were passed through a 100 μm strainer and centrifuged to remove collagenase solution. Single cell suspension was stained with viability dye on ice before performing flow cytometry. Intestinal tissue was then cut into 1-2 cm pieces and incubated in 25mL IEL solution [1x PBS w/ 2% FBS, 10mM HEPES buffer, 1% penicillin/streptomycin, 1% L-glutamine, plus 1mM EDTA (Sigma, E4884) and 1mM DTT (Sigma, D9779) added immediately before use] for 15 minutes at 37°C with vigorous shaking (250rpm). After centrifugation (450 g, 5 minutes) cells and tissue were resuspended in wash buffer (1x RPMI 1640 w/ 2% FBS, 10mM HEPES buffer, 1% penicillin/streptomycin, 1% L-glutamine) by vortexing. Epithelial and immune cells from the epithelial layer were removed by pouring suspension through a 100 μm strainer (ThermoFisher, 07-201-432). Remaining tissue was then incubated in 25mL collagenase and DNase I solution for 30 minutes at 37°C while shaking horizontally (250rpm) with ceramic beads (3 per sample). Lung tissue was placed in 5mL snap-cap tubes in 3mL of collagenase A and 1U/mL DNase I containing media, along with ceramic beads (3 per sample) and shaken horizontally at 250RPM for 45 minutes at 37°C. Digested samples were then passed through a 100 μm strainer and centrifuged to remove collagenase solution. Samples were then treated with 1x ACK to lyse red blood cells. Intestine and lung samples were washed by centrifugation in 40% Percoll™ (ThermoFisher) in 1x PBS to remove debris and enrich for lymphocytes prior to staining. All flow cytometry staining was done in Staining buffer on ice. For staining cytokine production, cells were incubated in Complete RPMI supplemented with 50ng/mL Phorbol 12-myristate 13-acetate (PMA) and 500ng/mL ionomycin with 1μg/mL brefeldin A and 2uM monensin to inhibit ER and Golgi transport for 3 hours at 37°C with 5% $CO_2$. For proenkephalin staining cells were additionally treated with 5 $\mu$M of a cell permeable proprotein convertase inhibitor (Millipore cat# 537076). Cells were stained for intracellular antigens using the CytoFix/CytoPerm kit (BD Biosciences) according to manufacturer instructions, adjusted for 96-well staining (100µl fix, 100µl stain, 200µl washes). Cell counts were performed using volumetric count. Samples were acquired on an Aurora (Cytek) and analyzed using FlowJo software (BD Biosciences).

**Transfection**

HEK293T cells were transfected with *Penk-P2A-eGFP* or *eGFP* in pcDNA3.1 plasmid backbone using Lipofectamine 3000 Transfection Reagent (Life technologies) according to manufacturer's instructions. Cells were analyzed by flow cytometry 48 hours after transfection.

**Antibodies**

The following antibodies were used for flow cytometry and microscopy experiments: CD25 (PC61), CD4 (GK1.5), CD62L (MEL-14), Siglec-F (E50-2440), TCR gamma/delta (GL3), CD90.2 (53-2.1), CD8 (53-6.7), CD90.1 (HIS51), and TCR V$\beta$3 (KJ25) were purchased from BD Biosciences; CD45 (30-F11), CD73 (TY/11.8), CD44 (IM7) Ly-6C (HK1.4), CD11b (M1/70), CD8$\alpha$ (53-6.7), CD19 (6D5), FcεRIα (MAR-1), CD64 (X54-5/7.1), IL-5 (TRFK5), CD3e (17A2), IFN$\gamma$ (XMG1.2), IL-2 (JES6-5H4), IL-10 (JES5-16E3), Tub$\beta$3 (Tuj1), TCR V$\beta$5.1/5.2 (MR9-4), TCR V$\beta$8.1/8.2 (KJ16-133.18), TCR V$\beta$11 (KT11), and Zombie NIR were purchased from Biolegend; Foxp3 (FJK-16s), Gr-1 (RB6-8C5), c-Kit (2B8), CD11c (N418), TCR$\beta$ (H57-597), IL-17A (eBio17B7), IL-22 (1H8PWSR), IL-4 (11B11), PENK (PA5-53021), TCR V$\beta$6 (RR4-7), TCR V$\beta$12 (MR11-1) and IL-13 (eBio13A) were purchased from Thermo Fisher; MHC Class II (I-A/I-E) (M5/114.15.2) was purchased from Tonbo Biosciences; rabbit polyclonal against DsRed was purchased form Takara; chicken polyclonal against GFP (#A10262) was purchased from Invitrogen; and cFos (rabbit monoclonal, 9F6) was purchased from Cell Signaling Technology.

**ELISA**

Ear skin tissue was snap frozen in liquid nitrogen. Thawed tissue was weighed and resuspended in Tissue Extraction Reagent I (Thermo Scientific) supplied with Complete EDTA-free Protease Inhibitor Cocktail (Sigma-Aldrich) at 0.1 g tissue/ml. Tissues were homogenized in Soft Tissue homogenizing tubes with a Bead Ruptor 24 (Biotage) [Output: 6, On time: 30 s, Off time: 5 s, Cycles: 4]. IL-23(p40) was measured using purified capture antibody (C15.6, 4 µg/ml, Biolegend), biotin conjugated detection antibody (C17.8; 1 µg/ml, Biolegend), and avidin-conjugated HRP (eBioscience). ELISAs were read at OD 450 on a Synergy HTX instrument (BioTek). Background obtained with the secondary antibody alone was subtracted.

**Histopathological analysis**

Tissue samples were fixed in 10% neutral buffered formalin, processed for hematoxylin and eosin staining by Histowiz, Inc. (histowiz.com).

**Single cell RNA-seq analysis**

*Preparation of reference genome*: The mm39 mouse genome assembly and NCBI RefSeq annotation information (GTF file) were downloaded from the UCSC Genome browser(*62-65*). In order to account for the presence of the *Penk*$^{Cre}$, *Foxp3*$^{Thy1.1}$, and *Gt(ROSA)26Sor*$^{LSL\text{-}YFP}$ targeted mutations, the corresponding sequences were inserted into the appropriate locations of the mm39 genome using the 'reform' script, creating the 'reformed mm39' genome. The GTF file was modified to appropriately extend the *Penk*, *Foxp3*, and *Gt(ROSA)26Sor* transcript and gene annotations, and to shift all other affected annotations, resulting in a 'reformed GTF' using a custom R script, relying on the 'GenomicRanges' and 'rtracklayer' packages(*66, 67*). The reformed mm39 and reformed GTF were used for smartseq2 RNA-seq alignment and analyses after generating a STAR genome index using STAR (version 2.7.3a) with the following command(*68*):

RNA:

STAR --runMode genomeGenerate --runThreadN 4 --genomeDir mm39_100_RNA --genomeFastaFiles mm39_reformed.fa --sjdbGTFfile mm39_reformed.gtf

*smartseq2 RNA-seq data processing*: Samples were processed and aligned using Trimmomatic (version 0.39), STAR (version 2.7.3a), and samtools (version 1.12), with the following steps, where *Sample* stands in for each well in the smartseq2 plates (including positive and negative control wells) (*68-70*):

- TrimmomaticPE *Sample*_R1.fastq.gz *Sample*_R2.fastq.gz -baseout *Sample*.fastq.gz ILLUMINACLIP:TruSeq3-PE.fa:2:30:10 LEADING:3 TRAILING:3 SLIDINGWINDOW:4:15 MINLEN:36
- STAR --runThreadN 6 --runMode alignReads --genomeLoad NoSharedMemory --readFilesCommand zcat --genomeDir mm39_100_RNA --readFilesIn *Sample*_1P.fastq.gz *Sample*_2P.fastq.gz --outFileNamePrefix *Sample* --outSAMtype BAM Unsorted --outBAMcompression 6 --outFilterMultimapNmax 1 --

outFilterMismatchNoverLmax 0.06 --outFilterMatchNminOverLread 0.35 --outFilterMatchNmin 30 --alignEndsType EndToEnd

- samtools sort -@ 4 -n -o *Sample*.bam *Sample*Aligned.out.bam
- samtools fixmate -@ 4 -rm *Sample*.bam *Sample*.fixmate.bam
- samtools sort -@ 4 -o *Sample*.resort.bam *Sample*.fixmate.bam
- samtools markdup -@ 4 -l 1500 -r -s *Sample*.resort.bam *Sample*.duprm.bam
- samtools index -@ 4 -b *Sample*.duprm.bam

This procedure resulted in retention of all uniquely aligning reads, with PCR duplicates removed, to be used for downstream analysis. Reads aligning to genes present in the reformed GTF were then counted using a custom R script relying on the 'GenomicAlignments', 'GenomicRanges', and 'GenomicFeatures' packages with default counting parameters(*66*).

*smartseq2 RNA-seq analysis:* Smartseq2 data were further processed and analyzed with a custom R script relying on the 'Seurat' (version 4) package (bioRxiv 2020.10.12.335331; doi: https://doi.org/10.1101/2020.10.12.335331). Counts for each plate were turned into a SeuratObject and normalized. Single cell containing wells with a number of detected genes (nFeature_RNA) fewer than 1.5x the highest number of genes detected in any negative control well were eliminated, as were wells containing numbers of detected genes (nFeature_RNA) greater than the mean number of genes detected in any positive control well (10 sorted cells). After this, the cells from the four individual wells were merged and analyzed together. First, the top 2000 variable genes were identified and scaled. A PCA was performed on these genes and the top 20 principal components were used to assign k-nearest neighbors, generate a shared nearest neighbor graph, and then optimize the modularity function to determine clusters, at resolution = 1.0(*71*). The shared nearest neighbor graph was used as input for the python-based algorithm 'Harmony' (600 iterations) in order to generate a two dimensional force directed layout for visualization(*72*). The 20 PCs were used as input for the python-based algorithm 'Palantir' in order to determine 'pseudotime' values(*73*), using a cell from cluster 0, which had the lowest per cell transcript counts, as the starting cell. Differential expression analysis for cluster 2, which was highly enriched for cells with a history of *Penk* expression, was performed using the *FindMarkers* function, with no fold change cutoff, and excluding genes expressed in fewer than 6.25% of cells (> 3 cells in the largest cluster). Significance testing was carried out with the Wilcoxon Rank Sum test using the Bonferroni correction, with an alpha of 0.1 for significantly differentially expressed. TCR activated and repressed genes were defined as genes which lost and gained expression in $T_{reg}$ cells ablated of the *Tcra* gene(*74*). The 'Seurat' function *AddModuleScore* was used to assign scores for the TCR activated and repressed genesets to each cell.

*Statistical analysis and data plotting*: All statistical tests were carried out in R using base R or the indicated packages, or by the indicated software or python algorithms. Plots were generated in R, using the ‘ggplot2’, ‘patchwork’, ‘RColorBrewer’, and ‘Seurat’ packages.

**Statistical analysis**

All statistical analyses (excluding RNA-seq, described above) were performed using GraphPad Prism 6 software. Differences between individual groups were analyzed for statistical significance using unpaired or paired two-tailed t-test, or one-way ANOVA. *, $p < 0.05$; **, $p < 0.01$; ***, $p < 0.001$. The number of mice used in each experiment to reach statistical significance was determined on the basis of preliminary data. No animals were excluded from the analyses. No methods of randomization were used to allocate animals into experimental groups. No blinding was used. Data met assumptions of statistical methods used and variance was similar between groups that were statistically compared.

### Supplementary Figures

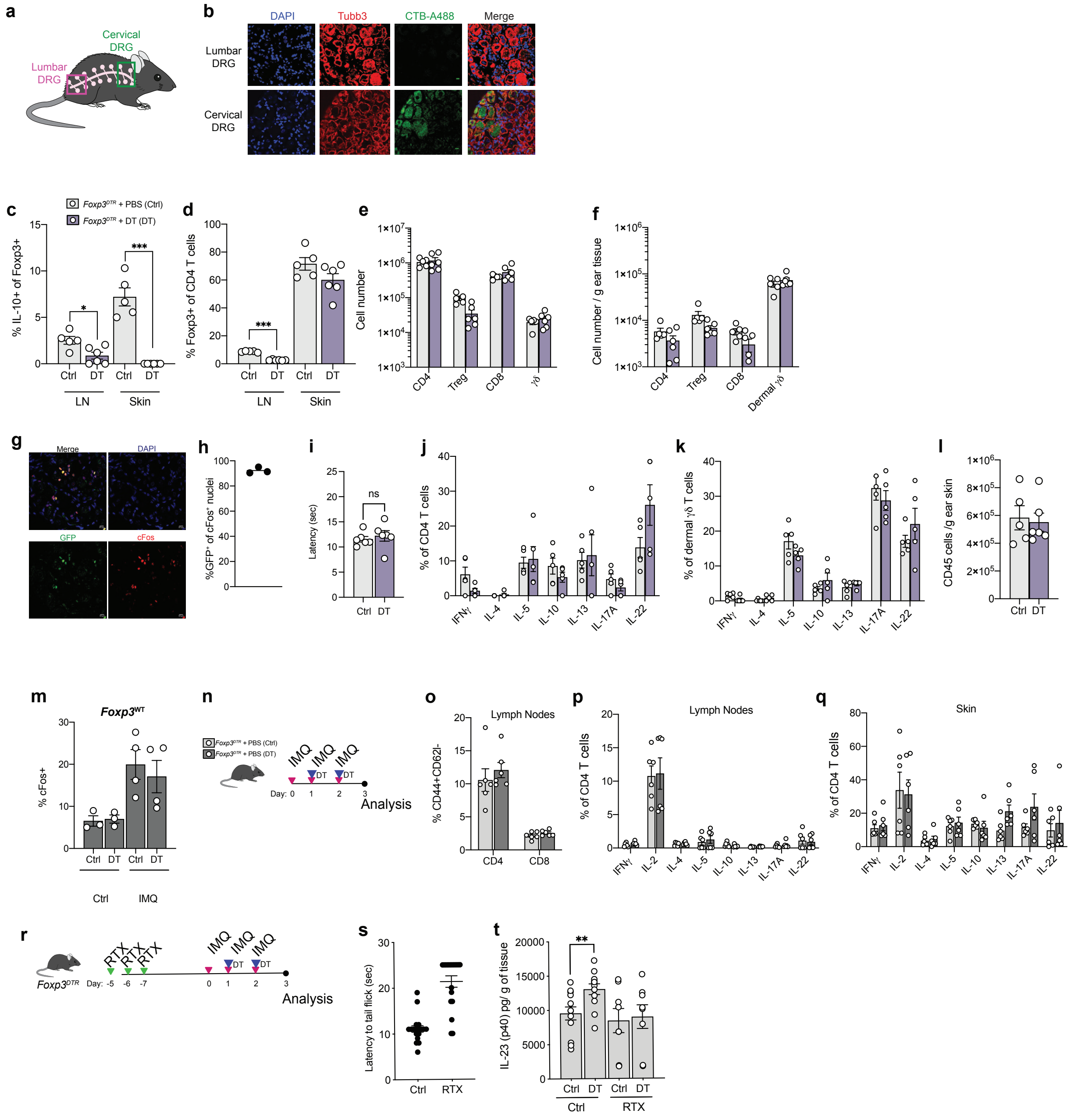

**Supplementary figure 1. Effects of $T_{reg}$ cell inactivation in *Foxp3$^{DTR}$* mice upon short-term DT treatment.**

**a-b** Retrograde labeling was performed by intradermal injection into the ear skin with CTB-A488. DRG tissues were taken 5 days after injection and analyzed by microscopy. **a,** Schematic of DRG isolated after retrograde tracing**. b,** Representative images of cervical and lumbar DRG.

**c-f,** *Foxp3$^{DTR}$* mice were treated with DT or PBS control. Cells were isolated from skin and LN 18 hours post DT treatment. **c,** Cells from indicated tissues were restimulated with PMA/ionomycin for 4 hours. IL-10 production was quantified by flow cytometry. **d,** $T_{reg}$ cell percentages of CD4 T cells isolated from LN and skin. **e-f,** Total cell number of indicated populations isolated from (g) LN and (h) skin.

**g-h,** DRG from *Fos-gfp* mice were isolated and analyzed by confocal microscopy. **g,** Representative confocal microscopy image of DRG sections stained for cFos (red), DAPI (blue) and GFP (green). **h,** Percentage of $GFP^+$ of nuclei stained by cFos antibody.

**i,** Latency to respond to thermal stimulus measured in female *Foxp$^{DTR}$* mice treated with DT or PBS control measured by hot plate assay 18 hours post DT treatment. Each dot represents response time per mouse.

**j-l,** *Foxp3$^{DTR}$* mice were treated with DT or PBS control. Cells were isolated from the skin 18 hours post DT treatment. **j-m,** Cytokine production was quantified by flow cytometry. Percentage of skin (l) CD4 T cells and (m) dermal $\gamma\delta$ T cells producing the indicated cytokines following 4 hours of PMA/ionomycin restimulation. **l,** Total number of $CD45^+$ cells isolated from skin.

**m,** *Foxp3$^{WT}$* mice were treated daily with topical IMQ or petroleum jelly control on ear skin (start: day 0). DT or PBS control was administered on day 1 and 2. Mice were analyzed on day 3. Percentage of $cFos^+$ nuclei of $Tub\beta3^+$ cells in DRG was quantified by confocal microscopy.

**n-q,** *Foxp3$^{DTR}$* mice were treated with IMQ for 3 consecutive days (start: day 0). DT or PBS control was administered on day 1 and 2. Tissues were collected on day 3. **n,** Schematic of treatments. **o,** Frequency of activated CD4 and CD8 T cells in the cervical lymph nodes following DT or control treatment. **p-q,** Percentage of CD4 T cells isolated from lymph nodes (r) and skin (s) producing the indicated cytokines following 4 hours of PMA/ionomycin restimulation.

**r-t,** *Foxp3$^{DTR}$* mice were treated with RTX or vehicle. One week post RTX treatment mice were treated daily with IMQ (start: day 0). DT or PBS control was administered on day 1 and 2. **r,** Schematic of treatments. **s,** One week following RTX or Ctrl treatment sensory denervation was tested by tail flick assay. **t,** Ear skin was collected on day 3 after IMQ treatment and IL-23 (p40) was measured by ELISA.

Each dot represents data from a mouse, bars show mean, error bars show SEM. *, $p<0.005$; **, $p<0.001$, ***, $p<0.0001$. P-values were calculated using an unpaired t-test.

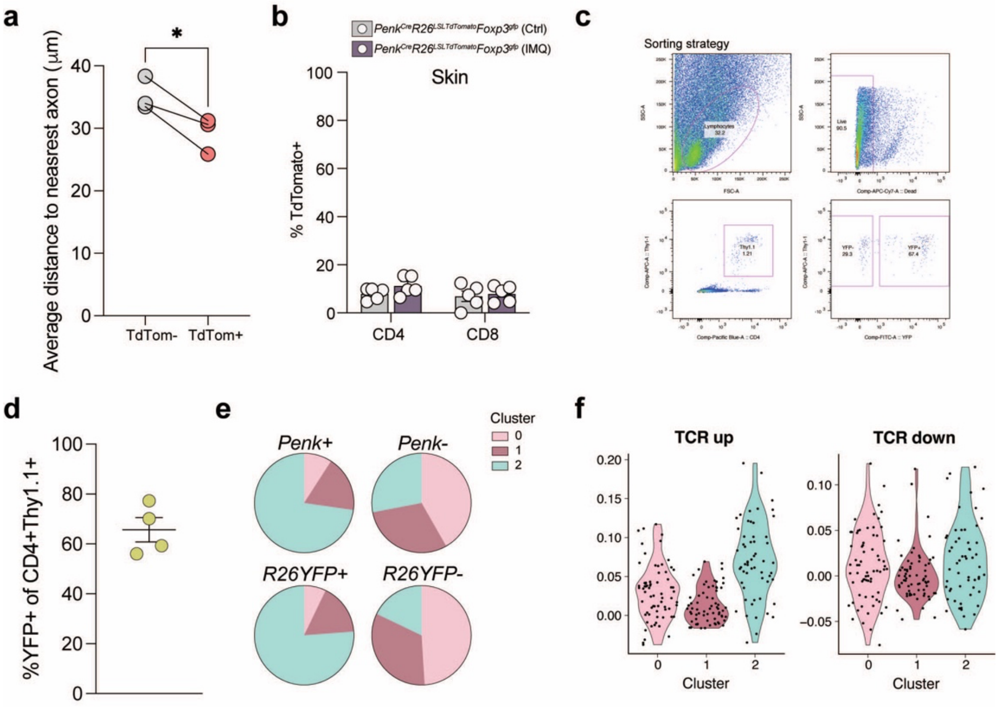


**Supplementary figure 2. A subset of skin Treg cells express Penk.**

**a,** Ear skin from *Penk*$^{Cre}$*R26*$^{LSL\text{-}TdTomato}$*Foxp3*$^{gfp}$ mice was analyzed by confocal microscopy. Average distance from tdTomato$^{+}$ and tdTomato$^{-}$ $T_{reg}$ cells to nearest axon per mouse. **b**, *Penk*$^{Cre}$*R26*$^{LSL\text{-}TdTomato}$*Foxp3*$^{gfp}$ were treated with petroleum jelly (ctrl) or IMQ for 3 consecutive days. Frequency of tdTomato$^{+}$ CD4 and CD8 T cells in skin. **c-f,** *Penk*$^{Cre}$*R26*$^{LSL\text{-}YFP}$*Foxp3*$^{Thy1.1}$ mice were treated with petroleum jelly (control) or IMQ for 3 consecutive days. $T_{reg}$ cells (Thy1.1$^{+}$CD4$^{+}$) were sorted from the ear skin of IMQ and control treated mice and processed for SMART-Seq2. **c**, Sorting strategy. **d**, Frequency of YFP$^{+}$ $T_{reg}$ cells in ear skin tissues from *Penk*$^{Cre}$*R26*$^{LSL\text{-}YFP}$*Foxp3*$^{Thy1.1}$ mice. **e**, Distribution of cells expressing or non-expressing specified transcripts across indicated clusters. **f**, Sequenced cells were scored for expression of sets of genes either TCR activated (TCR up) and TCR repressed (TCR down) in $T_{reg}$ cells. Violin plots depict genes per geneset score by each cluster. Each point represents a cell and violin contours represent distribution and spread. See *Methods* for details on genesets used.

Each dot represents data from a mouse, bars show mean, error bars show SEM. *, p<0.005. P-values were calculated using an paired t-test, or one-way ANOVA.

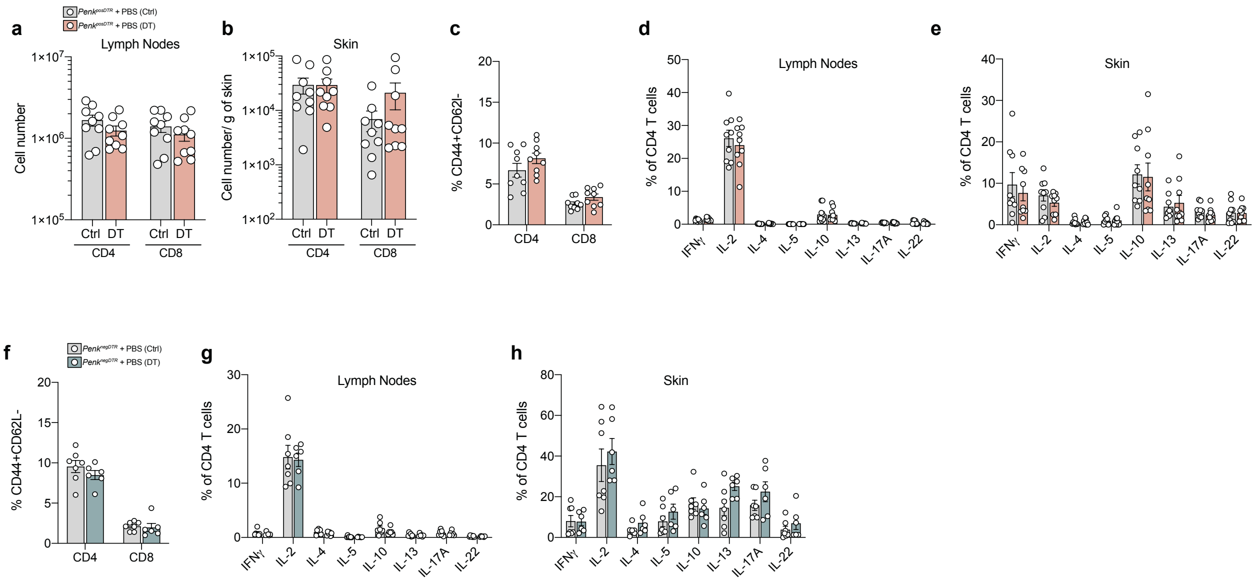


**Supplementary figure 3. Selective ablation of Penk-expressing and non-expressing $T_{reg}$ cells.**

**a-e,** *Tcrb*$^{-/-}$ mice were irradiated and reconstituted with a mix of bone marrow from *Tcrb*$^{-/-}$ and *R26*$^{iDTR}$*Penk*$^{Cre}$ at 4:1 ratio (referred to as *Penk*$^{pos}$*DTR*). 6 weeks post reconstitution *Penk*$^{pos}$*DTR* mice were treated with IMQ for 3 consecutive days (start: day 0). DT or PBS control was administered on day 1 and 2. Tissues were collected on day 3. **a-b,** Number of CD4 and CD8 T cells found in the cervical lymph nodes (a) and skin (b) following DT or PBS control treatment. **c,** Frequency of activated CD4 and CD8 T cells in the cervical lymph nodes following DT or control treatment in *Penk*$^{pos}$*DTR* mice. **d-e,** Percentage CD4 T cells isolated from lymph nodes (d) and skin (e) producing the indicated cytokines following 4 hours of PMA/ionomycin restimulation.

**f-h,** *Foxp3*$^{fl\text{-}DTR}$*Penk*$^{cre}$ (referred as *Penk*$^{neg}$*DTR)* mice were treated with IMQ for 3 consecutive days (start: day 0). DT or PBS control was administered on day 1 and 2. Tissues were collected on day 3. **f,** Frequency of activated CD4 and CD8 T cells in the cervical lymph nodes following DT or control treatment in *Penk*$^{neg}$*DTR* mice. **g-h,** Percentage CD4 T cells isolated from lymph nodes (g) and skin (h) producing the indicated cytokines following 4 hours of PMA/ionomycin restimulation.

Each dot represents data from a mouse, bars show mean, error bars show SEM.

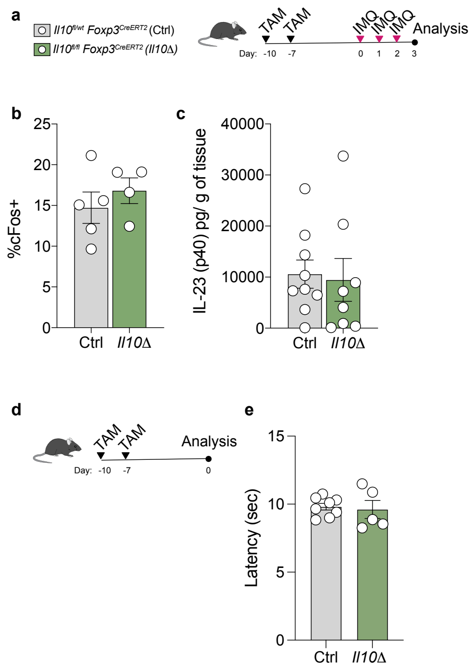


**Extended data figure 4. IL-10 ablation in $T_{reg}$ cells does not increase DRG activation or IL-23 production during early IMQ-induced inflammatory response.**

**a-c,** *Il10*$^{fl/wt}$*Foxp3*$^{creERT2}$ (Ctrl) and *Il10*$^{fl/fl}$*Foxp3*$^{creERT2}$ (*Il10Δ*) mice were treated with 2 doses of tamoxifen, 2 days apart. 7 days post tamoxifen administration Ctrl and *Il10Δ* mice were treated daily with IMQ for 3 days. **a,** Schematic of tamoxifen and IMQ administration. **b,** Percentage of cFos+ nuclei of Tubβ3 + cells in DRG. **c,** Concentration of IL-23 (p40)/ g of ear skin tissue quantified by ELISA.

**d-e**, Latency of response to thermal stimulus measured in male *Il10*$^{fl/wt}$*Foxp3*$^{creERT2}$ (Ctrl) and *Il10*$^{fl/fl}$*Foxp3*$^{creERT2}$ (*Il10Δ*) mice. Ctrl and *Il10Δ* mice were treated with 2 doses of tamoxifen, 3 days apart and analyzed 7 days post tamoxifen administration. **d,** Schematic of tamoxifen administration and time before analysis. **e** Response time in Ctrl and *Il10Δ* mice in hot plate assay.

Each dot represents data from a mouse, bars show mean, error bars show SEM.

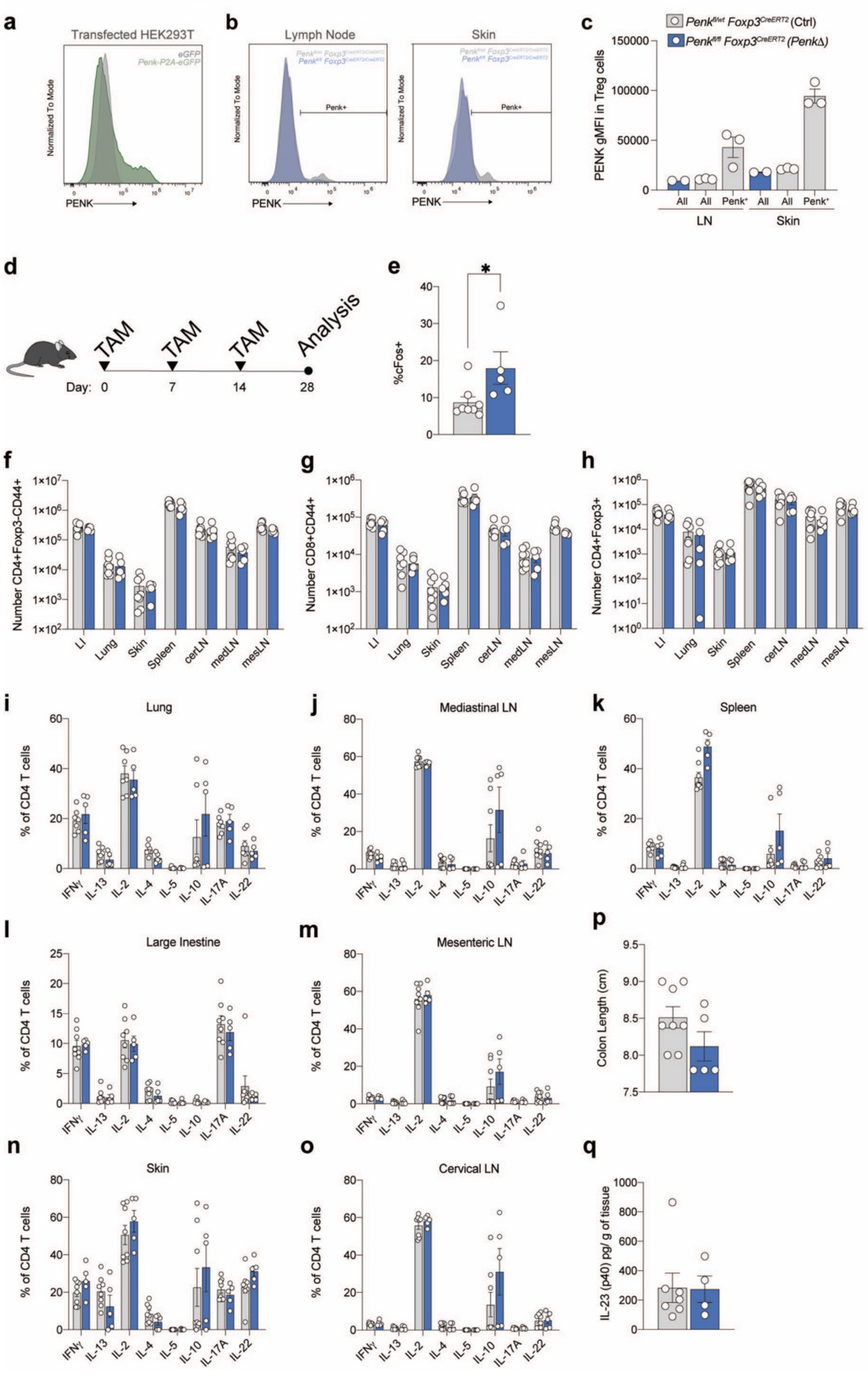


**Supplementary figure 5. Long term Penk ablation in $T_{reg}$ cells results in an increased DRG neuronal activity in the absence of immune activation and inflammation.**

**a,** HEK293T cells were transfected with *Penk-P2A-GFP* or *GFP* plasmid. 48 hours after transfection cells were stained for PENK by flow cytometry. Histogram shows representative PENK expression in GFP+ cells transfected with indicated plasmids**.**

**b-c,** *Penk*$^{fl/wt}$*Foxp3*$^{creERT2}$ (Ctrl) and *Penk*$^{fl/fl}$*Foxp3*$^{creERT2}$ (*PenkΔ*) were treated with 2 doses of tamoxifen, 2 days apart. 7 days post tamoxifen administration cells were isolated from lymph nodes and skin of Ctrl and *PenkΔ* mice, stained for PENK and analyzed by flow cytometry. **b,** Representative histogram of lymph node and skin from Ctrl and *PenkΔ* mice. **c**, Pooled mean fluorescence intensity (gMFI) from indicated populations in Ctrl and *PenkΔ* mice.

**d-q,** *Penk*$^{fl/wt}$*Foxp3*$^{creERT2}$ (Ctrl) and *Penk*$^{fl/fl}$*Foxp3*$^{creERT2}$ (*PenkΔ*) were treated with a weekly dose of tamoxifen for 28 days. 7 days post last tamoxifen administration cells were isolated from lymph nodes and skin of Ctrl and *PenkΔ* mice. **d**, Scheme of tamoxifen treatments and analysis time line. **e**, Percentage of cFos$^{+}$ nuclei of Tubβ3$^{+}$ cells in DRG. **f-h,** Number of activated CD4 (f), CD8 (g) and $T_{reg}$ cells (h) in the indicated tissues. **i-o,** Cytokine production by CD4 T cells isolated from (i) lung, (j) mediastinal lymph node, (k) spleen, (l) large intestine, (m) mesenteric lymph nodes, (n) skin, and (o) cervical lymph nodes, after 4 hours of PMA/Ionomycin restimulation. **p,** Colon length in Ctrl and *PenkΔ* mice. **q,** IL-23 (p40) concentration was measured by ELISA in Ctrl and *PenkΔ* mice.

Each dot represents data from a mouse, bars show mean, error bars show SEM. *, p<0.005. P-values were calculated using an unpaired t-test.

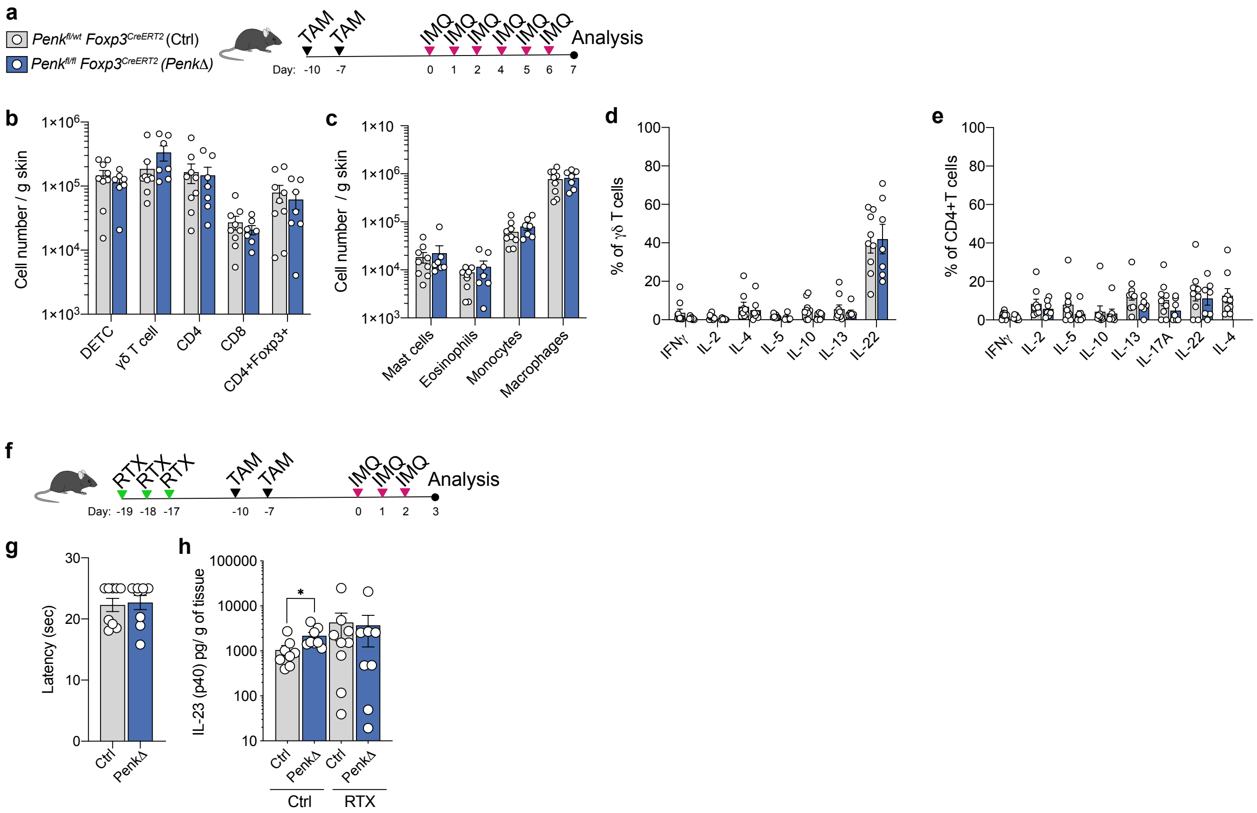


**Supplementary figure 6. Effects of $T_{reg}$ cell restricted *Penk* deficiency in psoriasiform skin inflammation.**

**a-e,** *Penk*$^{fl/wt}$*Foxp3*$^{creERT2}$ (Ctrl) and *Penk*$^{fl/fl}$*Foxp3*$^{creERT2}$ (*PenkΔ*) were treated with 2 doses of tamoxifen, 2 days apart. 7 days post tamoxifen administration Ctrl and *PenkΔ* mice were treated daily with IMQ for 7 days. **a,** Schematic of tamoxifen and IMQ administration. **b-c,** Number of indicated (b) T cell and (c) myeloid cell subsets in the skin of Ctrl and *PenkΔ* mice. **d-e,** Cytokine production by (d) γδ T cells and (e) CD4 T cells isolated from skin of Ctrl and *PenkΔ* mice after 4 hours of PMA/Ionomycin restimulation.

**f-h,** Ctrl and *PenkΔ* mice were treated with RTX or vehicle. **f,** Schematic of RTX, tamoxifen and IMQ administration. **g,** RTX treated Ctrl and *PenkΔ* mice were placed on a 58°C hot plate to measure latency of response to thermal stimulus 7 days after RTX treatment. Data shows response time to heat. **h,** One week post RTX or vehicle treatment mice were administered 2 doses of tamoxifen 2 days apart. 7 days following the last tamoxifen dose mice were treated daily with IMQ (start: day 0). Ear skin was collected on day 3 and IL-23 (p40) was measured by ELISA.

Each dot represents data from a mouse, bars show mean, error bars show SEM. *, $p<0.005$. P-values were calculated using an unpaired t-test.
